# Microglia Depletion Selectively Eliminates a Singular Form of Hippocampal Long-Term Potentiation

**DOI:** 10.1101/2022.07.29.501926

**Authors:** Jasmine Chavez, Aliza A. Le, Julian Quintanilla, Alex Mabou Tagne, Daniele Piomelli, Gary Lynch, Christine M. Gall

**Affiliations:** Departments of Anatomy and Neurobiology, University of California at Irvine, Irvine, California 92697, USA; Departments of Neurobiology and Behavior, University of California at Irvine, Irvine, California 92697, USA; Departments of Psychiatry and Human Behavior, University of California at Irvine, Irvine, California 92697, USA

**Author notes:** indicates equal contribution (Jasmine Chavez, Aliza A. Le, Julian Quintanilla). **Corresponding author:** Christine M. Gall, Gillespie Neuroscience Research Facility, 837 Health Science Road, Rm 3226, University of California, Irvine, CA 92697. **Author emails:** Jasmine Chavez, Julian Quintanilla, Aliza A. Le, Alex Mabou Tagne, Daniele Piomelli, Christine M. Gall, Gary Lynch.

**Keywords:** PLX5622, lateral perforant path, dentate gyrus, synaptic plasticity

## Abstract

There has been considerable recent interest in the possibility that microglia contribute to synaptic plasticity and some forms of learning. We report here that elimination of the cells in young adult male mice with a 7-12 day treatment with an antagonist (PLX5622) of the colony stimulating factor 1 receptor causes a profound but highly selective impairment to long-term potentiation (LTP) expressed by lateral perforant path (LPP) synapses with the dentate gyrus. Input/output functions and frequency facilitation to repetitive stimulation were not measurably affected. Direct infusion of PLX5622 into slices from naiive mice did not reduce the magnitude of LPP-LTP. Microglial depletion had no detectable effect on LTP in either the medial perforant path input to the dentate gyrus or the Schaffer-commissural projections between fields CA3 and CA1. We conclude that microglia discretely regulate the unusual form of LTP expressed by the LPP and thus exert region-specific effects on circuit function within hippocampus.

## Introduction

Microglia have historically been associated with removal of degenerating neuronal processes (Fu et al., 2014), but more recent work suggests that they participate in a diverse array of brain operations both during development and in adulthood (Erblich et al., 2011; Kierdorf and Prinz, 2017; Nelson et al., 2019; Andoh and Koyama, 2021; Dziabis and Bilbo, 2022). These broader contributions are typically thought to involve the elimination of connections either during pruning of excess synapses (Paolicelli et al., 2011; Kettenmann et al., 2013) or as a component of the response to insults, inflammation, or neurodegenerative disorders (Schafer et al., 2012; Kettenmann et al., 2013; Kierdorf and Prinz, 2017). Such functions fit well with the role of microglia as resident macrophages that release multiple bioactive agents in response to central and peripheral signaling molecules (Thomas, 1992; Kettenmann, 2007; Rivest, 2009). It also accords with the manner in which the cells continuously extend and retract thin branches (Bolton et al., 2022), a process thought to provide means for detecting changes in the local environment. It has been proposed that rapid scanning and sensing functions extend to events occurring during the normal range of synaptic activities and thus may be involved in different versions of memory-related plasticity (Morris et al., 2013). Related to this, genomic manipulations have implicated key microglial signaling systems in the regulation of long-term potentiation (LTP) (Maggi et al., 2011; Rogers et al., 2011; Pfeiffer et al., 2016; Zhou et al., 2019). Other studies have shown that such treatments or microglial depletion cause significant impairments to fear conditioning or motor learning (Parkhurst et al., 2013). Despite this progress, evidence that microglial elimination disrupts one or more LTP variants in forebrain is lacking.

Related to the above, multiple lines of evidence indicate that different projection systems utilize different forms of LTP. Early work on the hippocampus showed that enduring activity-induced potentiation in the Schaffer-commissural (SC) connections between fields CA3 and CA1 is both induced and expressed postsynaptically (Muller et al., 1988; Huganir and Nicoll, 2013; Lynch and Gall, 2013; Granger and Nicoll, 2014) whereas potentiation of the mossy fiber innervation of CA3 is generated presynaptically (Staubli, 1992; Castillo et al., 2002). The singular form of LTP present in the lateral perforant path (LPP) connections with the outer molecular layer of the dentate gyrus (DG) is initiated postsynaptically but expressed by changes in presynaptic release (Wang et al., 2016). These observations point to the possibility that regionally specific synaptic modification mechanisms respond in different ways to microglial signaling. If so, then the relationship of microglia to learning-related plasticity may be considerably more complex than typically envisioned. The studies reported here tested this possibility by evaluating the effect of pharmacological microglia ablation, using the colony stimulating factor 1 receptor (CSF1R) antagonist PLX5622, on LTP in the two perforant path inputs to the DG and the projections from field CA3 to CA1. The results show that microglial depletion severely impairs LPP-LTP without depressing potentiation in the other two pathways.

## Materials and Methods

Studies used male C57/BL6 mice (Charles River) at 2-4 months of age. Mice were group-housed (3-5 per cage) with access to food and water *ad libitum* and maintained on a 12 h light/dark cycle with lights on at 6:30 AM. Electrophysiological experiments were initiated from 8-10 AM. All procedures were conducted in accordance with NIH guidelines for the Care and Use of Laboratory Animals and Protocols and institutional approved protocols.

### PLX5622 treatment

To deplete brain microglia, mice were provided chow containing PLX5622 (1200 ppm) (Cayman Chemical Company, Ann Arbor, MI, USA), an antagonist of the CSF1R which is critical for microglial survival (Green et al., 2020; Liu et al., 2021). Poloxamine was included in the PLX5622 chow formulation to prevent recrystallization (Kuntsche et al., 2010). Animals were given control or PLX5622-containing chow and water *ad libitum* for 7-12 days; i.e., until euthanization. There were no differences between food consumption or body weights between the control and PLX groups over the treatment period.

### Slice preparation and recording procedures

Hippocampal slices were prepared for field recordings as previously published (Cox et al., 2019; Le et al., 2022; Quintanilla et al., 2022). For DG recordings, animals were anesthetized with isoflurane and euthanized by decapitation. Brains were rapidly removed and placed in ice-cold oxygenated (95% O_2_/ 5% CO_2_) high Mg^2+^ artificial cerebrospinal fluid (aCSF) containing (in mM): 87 NaCl, 26 NaHCO_3_, 25 glucose, 75 sucrose, 2.5 KCl, 1.25 NaH_2_PO_4_, 0.5 CaCl_2_, 7 MgCl_2_, (320-335mOsm). Horizontal slices (360μm) were then cut on the Leica vibrating tissue slicer (model, VT1000s, Leica). For CA1 recordings, transverse slices were prepared on a McIllwain chopper (360μm) and collected into ice-cold aCSF. Slices were immediately transferred to an interface recording chamber with a constant perfusion of oxygenated aCSF (31±1°C, 95% O_2_/ 5% CO_2_) containing the following (in mM): 124 NaCl, 3 KCl, 1.25 KH_2_PO_4_, 1.5 MgSO_4_, 26 NaHCO_3_, 2.5 CaCl_2_, and 10 dextrose at a rate of 60-70ml/h. Experiments were initiated ~1.5 h after slices were placed in the recording chamber. For all studies, stimulation was adjusted to elicit a field excitatory post-synaptic potential (fEPSP) of amplitude approximately 50% of the maximum population-spike free response. The fEPSPs were elicited using a bipolar stimulating electrode (twisted 65 μm nichrome wire) and recorded using a drawn glass pipette electrode containing 2M NaCl (2-3 MΩ). fEPSP slopes were collected using NACGather 2.0 (Theta Burst Corp., Irvine, CA, USA). For all slices, input-output curves were assessed, as were responses to a ten pulse 40 Hz stimulation train. Afterwards, stable baseline responses were collected for at least 20 min prior to induction of LTP as described below.

### Lateral Perforant Path (LPP)-DG

For the LPP-DG experiments, stimulating and recording electrodes were placed in the outer third of the DG molecular layer (internal blade). Proper electrode placement was confirmed using paired-pulse stimulation (40ms interval), which elicits response facilitation in the LPP but not medial perforant path (Christie and Abraham, 1994; Wang et al., 2016). For LPP-LTP experiments, stable baseline responses were collected for at least 20 min at 0.05 Hz and then potentiation was induced by applying 1s of 100 Hz, high-frequency stimulation (HFS) with pulse duration doubled and intensity increased x1.5 relative to baseline parameters; 0.05Hz stimulation resumed immediately after HFS and continued for 60 minutes.

### Medial Perforant Path (MPP) - DG

For MPP-DG analyses, recordings were performed in the presence of 1μM picrotoxin which was infused into the aCSF bath continuously from 40 min prior to experimentation (Amani et al., 2021). Stimulating and glass recording electrodes were placed in the middle third of the DG molecular layer. Electrode placement was verified by the presence of paired-pulse depression in response to a paired stimuli (40ms interval) as described (McNaughton, 1980; Christie and Abraham, 1994). After recording a stable 20 min baseline at 0.05 Hz, LTP was induced using three HFS trains (100Hz, 500ms each) with 20 seconds between trains at double duration and 1.5x intensity, relative to baseline. Recording responses to 0.05 Hz stimulation then resumed for an additional hour.

### Schaffer-Commissural (SC) - CA1

For the SC innervation of CA1 stratum radiatum, a stimulating electrode was placed in CA1c and the recording electrode was positioned in CA1b; both electrodes were located in stratum radiatum at the same distance from stratum pyramidale (Wang et al., 2018a). For LTP induction, a single five-burst train of theta burst stimulation (TBS) was used (4 pulses at 100Hz per burst, 200 ms between bursts) (Le et al., 2022) and 0.05 Hz stimulation resumed for one hour thereafter.

### PLX5622 Infusion

The CSF1R antagonist, PLX5622 (Cayman Chemical) was prepared as a stock solution by mixing 4.8:1 ratio (PLX:poloxamine) and dissolving the mixture in DMSO (10mM). The stock solution was then diluted in aCSF to achieve a final bath concentration of 1μM PLX (final DMSO concentration <0.01%). Vehicle solutions were made in the absence of PLX5622. After recording stable baseline responses, PLX5622 or vehicle was introduced into the bath via a secondary infusion line using a syringe pump (6mL/hr) for 60 minutes prior to induction of LTP, and for the duration of the experiment.

### Immunofluorescence

To verify microglial depletion with PLX5622 treatment, hippocampal slices were immersion fixed in 4% paraformaldehyde at the time of slice preparation or after electrophysiological recordings. The following day slices were sub-sectioned at a thickness of 25μm, mounted onto microscope slices, and processed for immunofluorescence as described (Rex et al., 2009; Seese et al., 2012; Wang et al., 2018a) using rabbit antisera to the microglial protein Iba1 (1:1000; Fujifilm Wako Pure Chemical Corp) and AlexaFluor Donkey anti-rabbit 594 (1:1000; Millipore). Slides were cover slipped with Vectashield containing DAPI (Vector laboratories).

To assess treatment effects on microglia, Iba1-immunoreactive (ir) cells were quantified within the full field of view (685 μm x 580 μm) for images collected at 20X objective magnification; this field extended from the DG granule cell layer (internal blade) to the hippocampal pyramidal cell layer at the CA1/subicular border (Figure 1); immunoreactive cells were counted from digital images.

**Figure 1:**
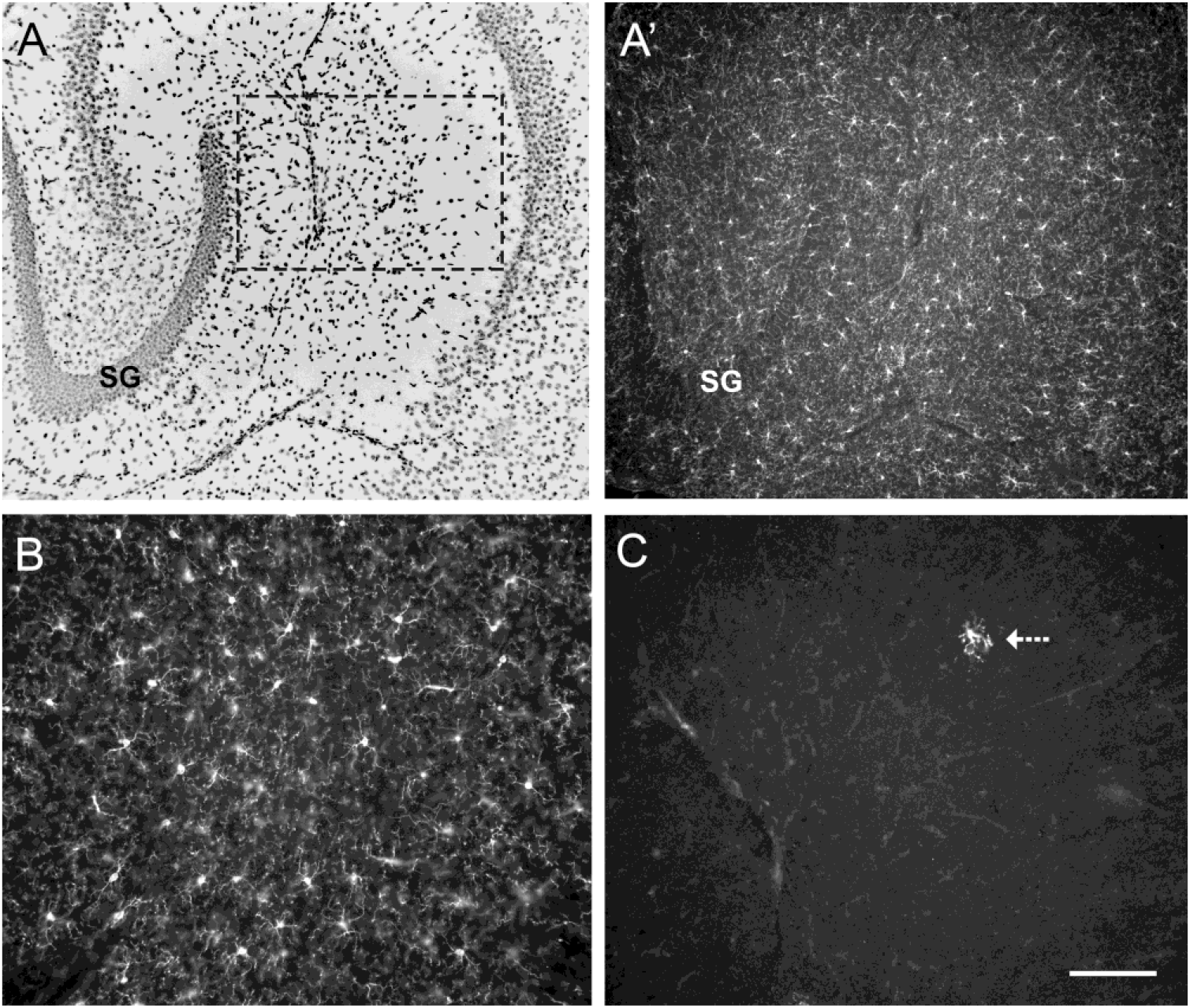
Treatment with PLX5622 depletes hippocampal microglia. **A**, **A’)** Images of the same hippocampal cross section viewed in different color channels showing DAPI-labeling of cellular nuclei (A, inverse contrast image) and Iba1 immunoreactivity (A’) in a slice from a vehicle-treated mouse fixed promptly after preparation (SG: dentate gyrus stratum granulosum; CA1 stratum pyramidale at right). Microglia appear white in A’-C. The dashed rectangle in A indicates the placement of the higher magnification images from which the Iba1-immunoreactive cells were quantified. **B, C)** Representative images of the quantification field showing Iba1 immunoreactivity in slices from vehicle- (B) and PLX5622- (C) treated mice: a severe depletion of Iba1-immunoreactive cells is evident in the PLX5622 case (arrow in C indicates the single immunolabeled cell in the field). Calibration bar in C indicates 200 μm for A and A’, 100 μm for B and C.

### Statistics

Results are presented as mean ± standard error of the mean (SEM) unless otherwise noted. Statistical significance (p<0.05) was determined using Excel and GraphPad Prism v6.0 (San Diego, CA). The significance of PLX5622 effects on immunolabeled cells was assessed using the 2-tailed t-test. For electrophysiological studies, the percent increase in response size following LTP induction was measured by comparing, on a slice by slice basis, the average fEPSP slope for the last 5 minutes of recordings to that for the 5 minutes prior to LTP inducing stimulation; significance was assessed using a 2-tailed unpaired t-test. 2-way repeated-measures (RM) ANOVA was employed for 40Hz train and theta burst response area comparisons. For all experiments, the “n” represents the number of hippocampal slices across at least 5 mice per group.

## Results

As described by others (Liu et al., 2004; Spangenberg et al., 2019), Iba1-ir microglial cells were evenly distributed across the hippocampal subfields in mice given control chow (VEH) (**Figs 1A, A’**); at higher magnification one could see the small Iba1-ir cells were highly ramified and distributed in a field of fine immunolabeled processes (**Fig 1B**). In accord with earlier reports (Liu et al., 2004; Huang et al., 2018; Spangenberg et al., 2019; Basilico et al., 2022), treatment with chow containing PLX5622 (PLX) caused a dramatic reduction in the number of microglia (**Fig 1C**) in all hippocampal subfields within 7 days and this depletion was sustained as animals continued on the diet. Quantification of Iba1-ir cells within a sample field extending across the dorsal blade of the DG molecular layer and the contiguous apical dendritic field of CA1 confirmed the presence of a near complete depletion with treatment (PLX: 3.48±2.03, VEH: 79.3±7.12; mean±standard deviation for a representative set of slices from 5 PLX and 4 VEH mice; p=7.39 × 10^-8^; 2-tailed unpaired t-test) (**Fig 1C**).

### Effects of microglial depletion on perforant pathway physiology

There were no evident effects of the treatment on baseline fEPSPs evoked by single pulse stimulation of the LPP and recorded in the outer molecular layer of the DG. Input/output curves (fiber volley vs. amplitude of the response) were superimposable in the vehicle vs. PLX groups (**Fig 2A**), indicating that microglial depletion did not affect the relationship between number of axons activated and the resultant magnitude of dendritic depolarization. LPP-DG synapses exhibit an atypical response to short trains of gamma frequency (40Hz) stimulation in which the fEPSP facilitates for the first two or three pulses and then become progressively more depressed as the train continues. This pattern was intact in the PLX treatment (F_9, 135_=0.37, p=0.95; PLX n=10, Veh n=7, two-way RM ANOVA) (**Fig 2B**). In all, a massive reduction in the microglia population did not significantly affect complex synaptic operations in the DG. Despite this, there was a dramatic change in LPP-LTP elicited by a single, one second long train of 100Hz stimulation. The first response after the HFS train was enhanced in the PLX group to about the same level as in vehicle controls but potentiation rapidly decayed to near baseline values (PLX: 115.8±6.1%, Veh: 147.7±6.8%, two-tailed unpaired t-test t_16_=3.51, p=0.0029; PLX n=9, Veh n=9) (**Fig 2C, D**). These findings indicate that microglia have surprisingly discrete effects in the DG and that these largely target the mechanisms underlying the production of lasting changes in synaptic strength.

**FIGURE 2:**
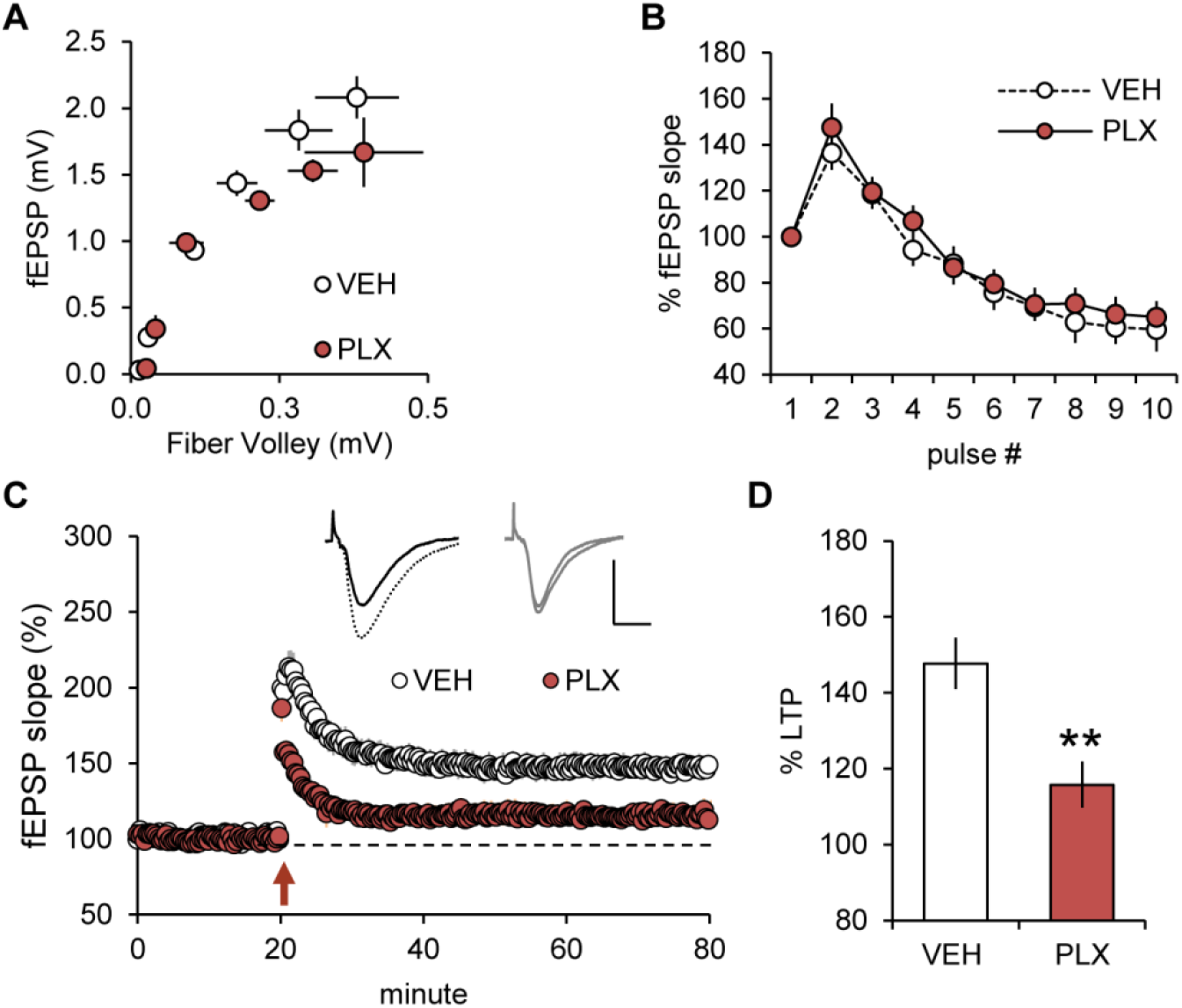
Lateral perforant path (LPP) LTP is markedly impaired in microglia depleted mice. Acute hippocampal slices from mice fed either control (VEH) or PLX5622- (PLX) chow for 7-12 days were assessed for treatment effects on LPP transmission and LTP. **A**) Input-output curves from the outer molecular layer of the DG for VEH and PLX treated animals; VEH n=8, PLX n=5. **B)** The response profile to a ten-pulse, 40 Hz train of stimulation was comparable between VEH and PLX groups, both showing the typical response facilitation followed by depression (F_9,135_=0.3653, p=0.9497, VEH n=7; PLX n=10; two-way RM ANOVA). **C**) After recording stable baseline responses, a single 100 Hz HFS train was used to induce LTP (at arrow). Both VEH and PLX groups exhibited an initial short-term potentiation and decline to a stable potentiated response, but the magnitude of potentiation was markedly reduced in the PLX group relative to VEH-controls. Representative traces show responses before (solid) and after (dashed) induction of LTP. Scale bars: x=1 mV, y=10 ms. **D)** Plot of the mean potentiation at 55-60 min post-induction showed mice treated with PLX had significantly reduced LPP-DG LTP relative to VEH mice (**p=0.0029, VEH n=9; PLX n=9; two-tailed unpaired t-test).

Next, we examined the effects of microglial depletion on responses elicited in the middle molecular layer of the DG by stimulation of the MPP. The fEPSPs were not discernably different in the vehicle vs. PLX cases and these groups had comparable input/output curves (**Fig 3A**). The MPP belongs to a category of synapses in which repetitive stimulation produces a strong within-train reduction of synaptic responses (McNaughton, 1980; Christie and Abraham, 1994). MPP-evoked responses, in both vehicle and PLX groups, exhibited similar degrees of depression across a ten pulse 40Hz train (**Fig 3B**) (F_9, 108_=1.09, p=0.38; Veh n=5, PLX n=9, two-way RM ANOVA). Finally, and in marked contrast to the results obtained for LPP-LTP, microglial depletion did not produce a significant change in MPP-LTP (t_11_=1.144 p=0.277, PLX n=7, Veh n=6, two-tailed unpaired t-test) (**Fig 3C**). The percent potentiation recorded 55-60 minutes after delivery of HFS was 162.2±9.7% for depleted slices and 148.2±6.9% for vehicle slices. This result indicates that microglia in the DG may target a cellular process that is essential for LPP-LTP but not involved in the production of MPP-LTP.

**Figure 3:**
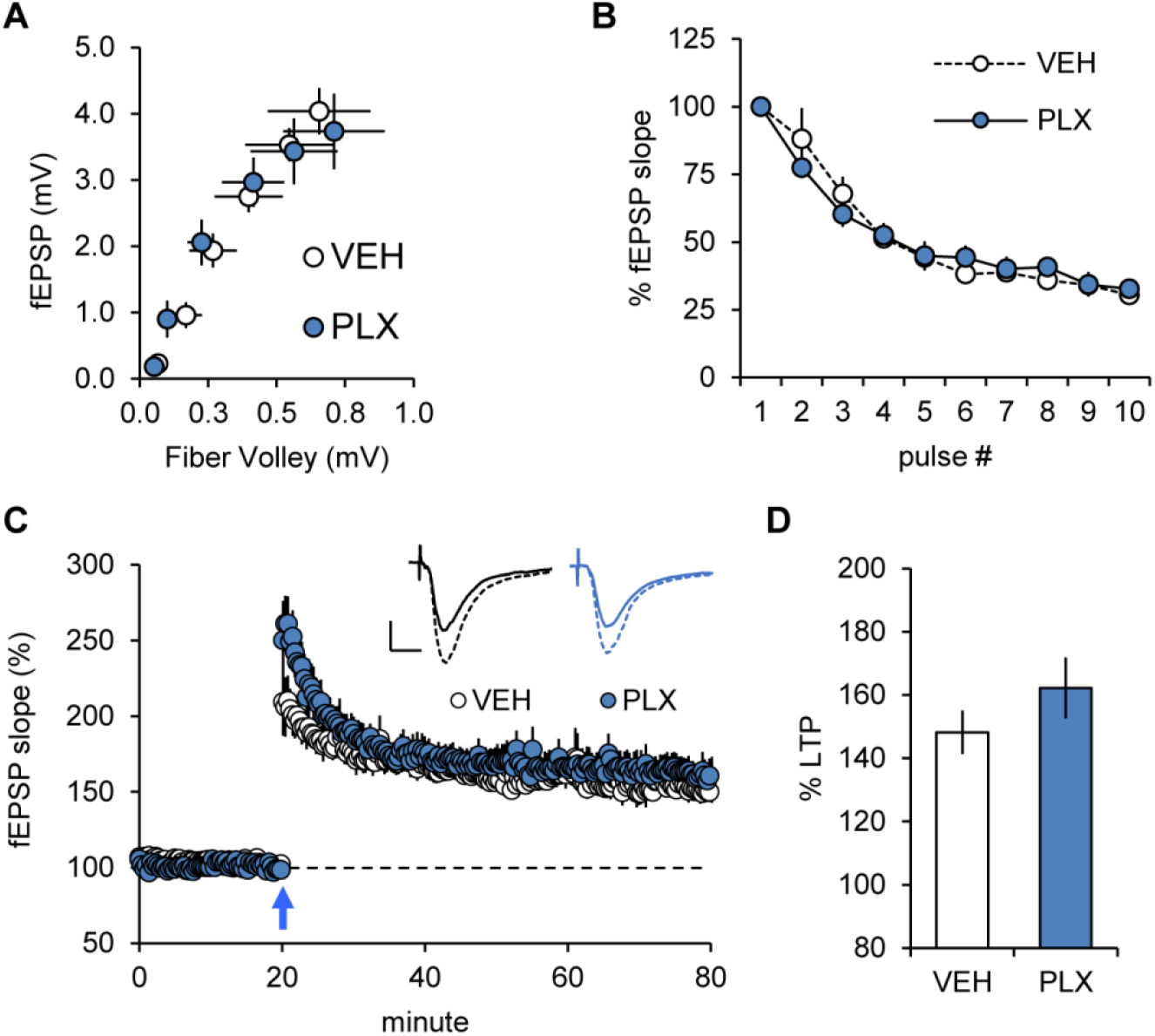
Microglial depletion did not affect medial perforant path (MPP) responses. **A)** Input/output curves for the MPP were comparable in slices from VEH- and PLX5622- (PLX) chow treated mice. **B)** Plot shows MPP-fEPSP responses to a ten pulse, 40 Hz train were comparable in VEH and PLX mice: in both groups responses were depressed by about 30% over the course of the train (F_9,108_= 1.09, p= 0.379; VEH n=5, PLX n=9; two-way RM ANOVA). **C)** After recording stable baseline responses for 20 min, MPP potentiation was induced using three, 500ms high frequency stimulation trains (100Hz each) separated by 20 sec (at arrow). Both VEH and PLX treated animals exhibited a marked potentiation. Scale bars: x=1 mV, y=10 ms. **D)** Plot of the mean fEPSP response recorded 55-60 min post-induction showed no difference in the degree of potentiation in slices from VEH and PLX mice (p=0.2768; VEH n=7, PLX n=6; two-tailed unpaired t-test).

### Microglial elimination and LTP in field CA1

Synaptic responses elicited in the proximal apical dendrites of field CA1b by single pulse stimulation of the SC projections from CA3 were not significantly changed by microglial depletion. As was the case for the DG, input/output curves were nearly identical in the vehicle and PLX groups (**Fig 4A**). The CA3-CA1, SC synapses exhibit a conventional frequency facilitation effect in response to short (10 pulse) trains of 40Hz stimulation. This profile was intact in the microglial elimination cases (F_9,99_= 0.30, p=0.97; two-way RM ANOVA) (**Fig 4B**). The size (area) of the response to individual theta bursts was not measurably changed by microglial depletion (F_4,52_=0.081, p=0.99) (**Fig 4C**). Applying a 5 TBS trains to the SC projections produced a rapid potentiation of the responses in CA1 that decayed over 10-15 minutes before stabilizing at about 50% above baseline levels in both vehicle and PLX groups. The lack of treatment effect was borne out by comparison of the magnitude of LTP as assessed at 55-60 minutes post TBS (PLX: 140.8±3.2%, Veh: 148.0±5.2%; p=0.24, 2-tailed unpaired t-test) (**Fig 4D**).

**FIGURE 4:**
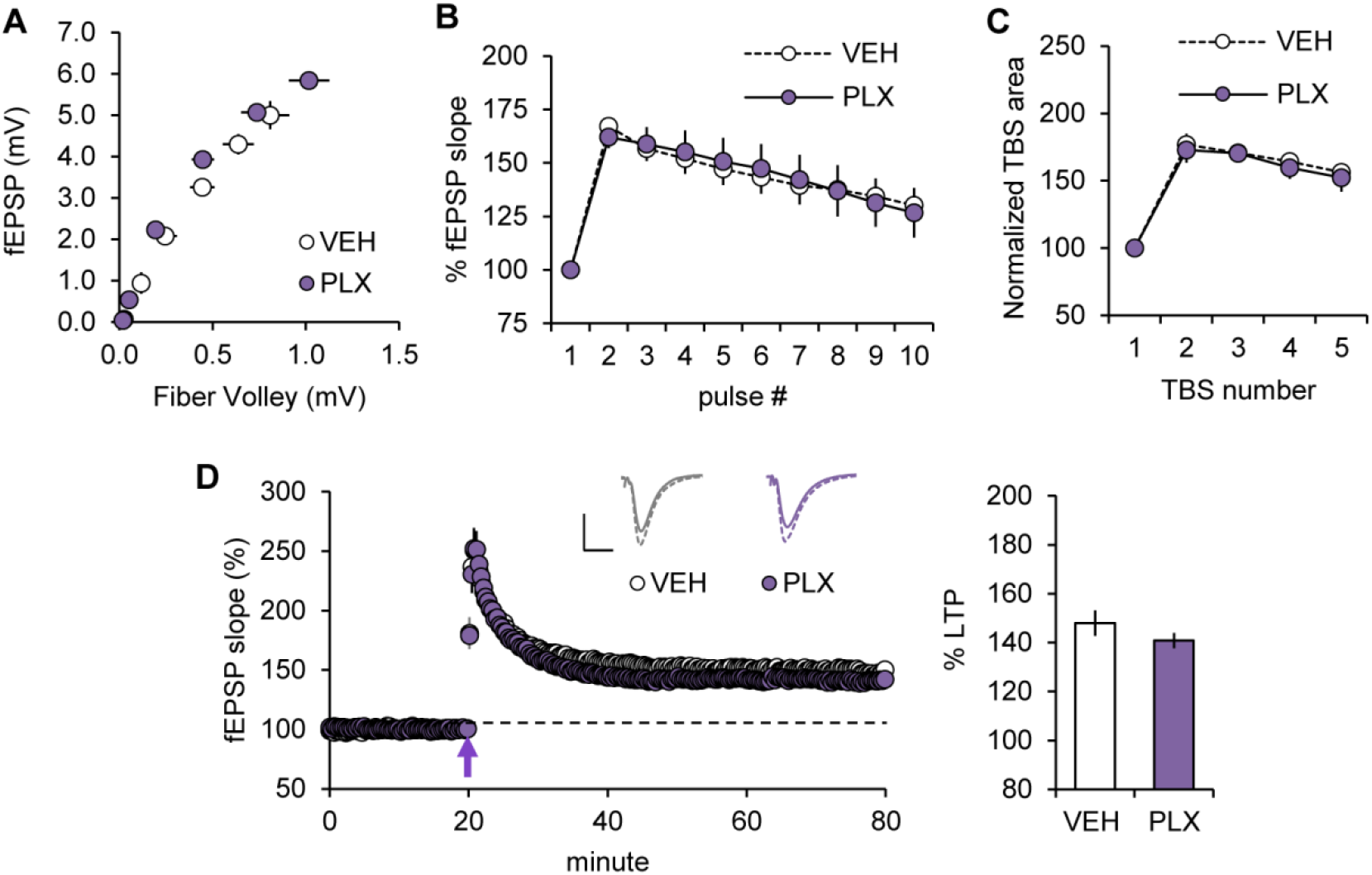
PLX5622 treatment did not influence field CA1, SC LTP. **A)** SC input/output curves were comparable between treatment groups. **B)** Plot shows SC fEPSP responses to a 10 pulse, 40 Hz train were comparable in VEH and PLX mice (F_9,99_= 0.3022, p=0.9725; VEH n=6, PLX5622 n=7; two-way RM ANOVA). **C)** There was no reliable effect of PLX on the area of the response to individual theta bursts used to induce LTP (F_4,52_=0.0812, p=0.988; VEH n=8, PLX n=7). **D)** A single train of 5 theta bursts elicited comparable potentiation in slices from VEH- and PLX-mice. Representative fEPSP traces collected before (solid line) and after (dotted line) LTP are shown for the two groups. The bar graph (right) summarizes the normalized slopes at 55-60 post TBS (p=0.2365; VEH n=7, PLX n=9; two-tailed unpaired t-test). Scale bars: x=1 mV, y=10 ms.

### Effects of acute infusion with PLX5622

We next assessed the effects of acute administration of the compound at a concentration that blocks CSF1R. Further analyses on the LPP-LTP were conducted because of the marked effect of microglial depletion on plasticity in this system. Infusion of the antagonist for 60-90 minutes had no evident effect on the waveform of LPP fEPSPs and did not reduce the magnitude of LTP (two-tailed unpaired t-test t_14_=0.0277 p=0.9783; PLX n=8, Veh n=8) (**Fig 5A,B**). Percent potentiation at 60 minutes post HFS was 156.0±11.8% for vehicle- and 155.5±14.5% for PLX-treated slices.

**FIGURE 5:**
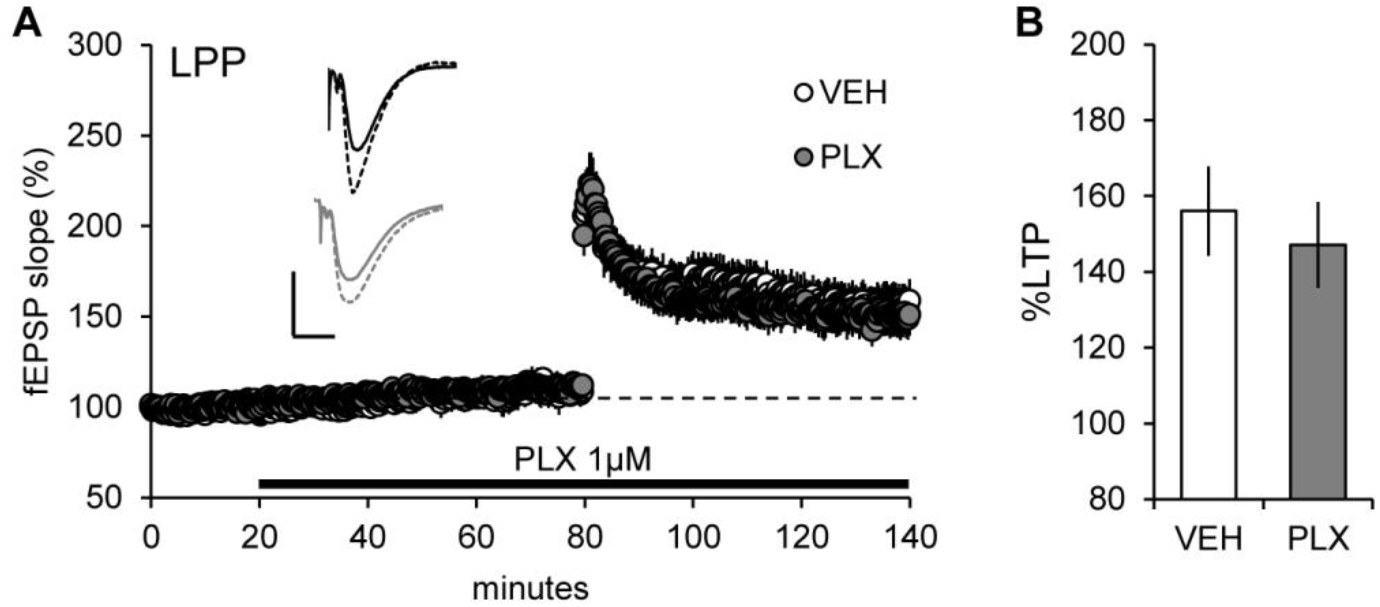
Acute infusion of PLX5622 does not impair LPP LTP. LPP fEPSP responses were collected from slices from naiive mice and were infused with VEH or 1μM of PLX5622 (solid bar). **A)** After infusion of VEH or PLX for 1 hour, a HFS (1 sec, 100Hz) was applied to induce LPP-LTP; the two groups exhibited similar levels of potentiation. Scale bars: x=1 mV, y=10 ms. **B)** Plot of the mean percent potentiation at 55-60 min after HFS shows comparable LPP-LTP in the Veh and PLX groups (p=0.590; VEH n=11, PLX n=8; two-tailed unpaired t-test).

## Discussion

The above results describe a surprisingly localized and selective effect of microglial depletion on synaptic plasticity in the adult hippocampus. The cellular events responsible for transferring synapses into their potentiated state differ between the systems exhibiting PLX5622 associated impairment and those unaffected. Regarding the former, LTP in the LPP-DG contacts is initiated in the target dendrites by on-demand synthesis of the endocannabinoid 2-arachidonoylglycerol (2-AG) which diffuses to LPP terminals where it binds to cannabinoid receptor 1 (CB_1_R) (Wang et al., 2016). These receptors typical depress neurotransmitter release via effects on presynaptic calcium channels and by phosphorylating the vesicular protein Munc18-1 (Schmitz et al., 2016). Although this mechanism is present in LPP terminals, there is a lower threshold CB_1_R cascade leading to focal adhesion kinase (FAK), one of two tyrosine kinases used by integrins to regulate the actin cytoskeleton. Available evidence indicates that the CB_1_R-FAK system, working on coordination with presynaptic integrins, is necessary for the production of enduring activity-driven increases in evoked transmitter release, which is the means of LPP-LTP expression (Wang et al., 2018b). The present results strongly suggest that microglia exert a necessary and positive influence on one or more elements in this complex series of events. A first step in developing a more specific hypothesis as to microglial involvement would entail distinguishing between acute, perhaps activity driven, contributions versus a role that is more constitutive (i.e., in which microglia are needed to maintain the system). Studies on synaptic plasticity in the spinal cord have produced evidence for an active system in which high levels of afferent input causes microglia to release a neurotrophin thereby causing a lasting increase in synaptic potency; these effects are blocked by antibodies against CSF1R (Zhou et al., 2019). We found that acutely blocking CSF1R has no effect on LPP-LTP but activity-induced signaling involving other messengers remains a distinct possibility. Microglia were not needed to maintain the integrity of LPP-DG synaptic operations: input/output curves and synaptic responses appeared normal in the microglia depleted mice as did the complex response profile elicited by a 40 Hz gamma train. Understanding the mode of action for microglia in LPP-LTP will likely require identification of factors released by the cells that interact with the pre- and/or post-synaptic components of this exotic form of plasticity at this particular contact.

In contrast to the LPP, marked reduction in the microglia population had little if any effect on synaptic potentiation in the MPP projections to the DG or in the SC connections between fields CA3 and CA1. Surprisingly little is known about the substrates for MPP-LTP but it is clear that these do not include the endocannabinoid signaling that is critical to LPP-LTP (Wang et al., 2016). It should also be noted that the present MPP results establish that microglial depletion does not cause global disturbances to the DG granule cells. CA1-LTP also involves different synaptic mechanisms than those utilized by the LPP. The CA1 potentiation effect is both induced and expressed postsynaptically and does not require endocannabinoid messengers (Lynch et al., 2007; Lynch and Gall, 2013; Granger and Nicoll, 2014; Wang et al., 2016). It is integrin dependent but unlike the LPP the pertinent adhesion dimers are located on the dendritic spine and belong to the large family that recognizes a consensus RGD sequence in the matrix binding ligands (Kramár and Lynch, 2003; Kramár et al., 2006; Babayan et al., 2012). The presynaptic integrins that support LPP-LTP do not belong to this group. It is thus possible that the loss of microglial release of proteinases (del Zoppo et al., 2007; Könnecke and Bechmann, 2013; Crapser et al., 2021) that generate matrix-derived ligands for presynaptic integrins could result in the disruption of LTP in the LPP but not other sites within the hippocampus.

The present studies addressed the contributions of microglia to complex synaptic physiology and plasticity in young adult mice. There are reasons to suspect that the results, at least for LTP, would be different at both younger and later ages. The potentiation effect as found in CA1 undergoes substantial changes from its initial appearance in the second postnatal week to the onset of puberty (Kramár and Lynch, 2003; Kramár et al., 2012) and it was recently shown that further adjustments occur during the transition into young adulthood (Le et al., 2022). Declines in CA1-LTP are evident by the beginning of middle-age in CA1 (Rex et al., 2005) and even earlier in the LPP (Amani et al., 2021). The functions of microglia also change over life with a prominent role in synaptic pruning during postnatal development and central contributions to pathology in the aged brain (Crapser et al., 2021). Interactions between the age-related changes to plasticity and those for microglial operations are not unlikely. It is interesting in this regard that a recent and detailed report described losses in dendritic spines and decreases in the frequency of spontaneous release events after microglial depletion in 5-week old mice (Basilico et al., 2022). That study also found a modest increase in CA1-LTP induced with high frequency stimulation of afferents from CA3. Taken together, these results and those from the present study point to a possible puberty-related narrowing of microglial role in regulating excitatory transmission. It will be of interest to directly test the point in future studies.

Finally, prior studies showed that transient silencing of the LPP blocks the ability of mice to encode any of the three primary elements – identity, location, and temporal order of events – of unsupervised episodic memory (Cox et al., 2019). Other work has implicated the pathway in the rapid acquisition of rewarded cues in animals that had past experience with the training paradigm (Wang et al., 2016). The results reported here therefore lead to the intriguing prediction that two essential forms of everyday learning are dependent on the normal functioning of microglia in a restricted locus within hippocampus. This would provide one route whereby the various peripheral factors that influence microglial functioning could interfere with hippocampal contributions to memory formation.

## FUNDING AND ACKNOWLEDGEMENTS

This research was supported by the National Institutes of Drug Abuse award P50 DA44118 to D.P. and R01HD101642-02 to C.M.G. and G.L. We thank Ms. Yue Qin Yao for assistance with immunofluorescence.

